# Function-guided protein design by deep manifold sampling

**DOI:** 10.1101/2021.12.22.473759

**Authors:** Vladimir Gligorijević, Daniel Berenberg, Stephen Ra, Andrew Watkins, Simon Kelow, Kyunghyun Cho, Richard Bonneau

## Abstract

Protein design is challenging because it requires searching through a vast combinatorial space that is only sparsely functional. Self-supervised learning approaches offer the potential to navigate through this space more effectively and thereby accelerate protein engineering. We introduce a sequence denoising autoencoder (DAE) that learns the manifold of protein sequences from a large amount of potentially unlabelled proteins. This DAE is combined with a function predictor that guides sampling towards sequences with higher levels of desired functions. We train the sequence DAE on more than 20M unlabeled protein sequences spanning many evolutionarily diverse protein families and train the function predictor on approximately 0.5M sequences with known function labels. At test time, we sample from the model by iteratively denoising a sequence while exploiting the gradients from the function predictor. We present a few preliminary case studies of protein design that demonstrate the effectiveness of this proposed approach, which we refer to as “deep manifold sampling”, including metal binding site addition, function-preserving diversification, and global fold change.

## 1 Introduction

Protein design has led to remarkable results in past decades in synthetic biology, agriculture, medicine, and nanotechnology, including the development of new enzymes, peptides and biosensors [1]. However, sequence space is large, discrete, and sparsely functional [2], where only a small fraction of sequences may fold into stable structural conformations. Taken together, these considerations present important challenges for automated and efficient exploration of design space.

Building on previous works on representation learning from large-scale protein sequence data [3, 4, 5, 6, 7, 8], we introduce a novel generative model-based approach called “deep manifold sampling” for accelerating function-guided protein design to explore sequence space more effectively. By combining a sequence denoising autoencoder (DAE) with a function classifier trained on roughly 0.5M sequences with known function annotations from the Swiss-Prot database [9], our deep manifold sampler is capable of generating diverse sequences of variable length with desired functions. Moreover, we conjecture that by using a non-autoregressive approach, our deep manifold sampler is able to perform more effective sampling than previous autoregressive models.

## 2 Related Work

Recent work has demonstrated success in learning semantically-rich representations of proteins that encapsulate both biophysical and evolutionary properties. In particular, language models (LM) using bi-directional long short-term memory (LSTM) [10] and attention-based [5] architectures and trained on protein sequences have yielded useful representations for many downstream tasks, including secondary structure and contact map predictions [5], structural comparison [10], remote homology detection [7], protein engineering and fitness landscape inference [3], and function prediction [11].

Other studies have focused on generative modeling for producing realistic protein structures — for example, using Generative Adversarial Networks (GAN) for creating pairwise distance maps [12] and variational autoencoders (VAE) for 3D coordinate generation of protein backbones [13] — and designing new sequences. One advantage in formulating a design problem with sequences has traditionally been the relative availability of data as compared to experimentally-determined structures. Balakrishnan et al. [14] used graphical models trained on multiple sequence alignments (MSA) to sample new sequences. More recently, VAEs [15, 16] have been used for designing novel enzymes and T-cell receptor proteins, obviating the need for MSA, but they have been largely limited to a single family of proteins. Additionally, a few generative models have been proposed for conditional design [17, 6, 8]: Greener *et al*. [17] use a VAE conditioned on structural features for generating sequences with metal binding sites, Madani *et al*. [6] use a conditional LM for sampling proteins, where each amino acid is sampled sequentially, and Shin *et al*. [8] use an autoregressive model to generate a nanobody sequence library with high expression levels.

Motivated by conditional design as translation task, we develop an approach for generating protein sequences with desired functions, where sequences are translated from an input sequence to an output sequence with higher property values. Our deep manifold sampling approach uses a denoising autoencoder (DAE), a self-supervised model that implicitly estimates the structure of the data-generating density by denoising stochastically corrupted training examples [18, 19, 20, 21]. We use a Markov chain Monte Carlo (MCMC) process [21] to sample from the density function learned by the encoder. We corrupt this sample and repeat the procedure above to produce a chain of samples from the DAE.

We also consider known issues with autoregressive models including decoding latency [22, 23], difficulty of parallelization at inference time [24, 25, 26], and exposure bias at test-time generation [27, 28]. By using a non-autoregressive modeling strategy, our deep manifold sampler is capable of predicting multiple mutations — including insertions and deletions — at different positions in a given sequence resulting in sequences of varying lengths. We conjecture that in doing so, our manifold sampler enables effective exploration of the overall fitness landscape of properties, resulting in diverse protein designs with desirable properties.

## 3 Methods

We propose to learn a protein sequence manifold by training a DAE on a large database of observed sequences spanning multiple protein families. Moreover, to ensure that generated sequences satisfy a set of desired functional constraints, we combine a protein function classifier with the DAE to guide sampling.

### 3.1 A sequence denoising autoencoder

Our goal is to generate a diverse set of protein sequences that exhibit a high level of a desired function using a sequence DAE [29]. We want to map an input sequence 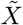 to a target sequence *X*, where *X* always has a higher level of some desired protein function than 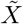. We formulate this task as a language modeling problem, where we model the joint probability of the tokens of a target protein sequence *X* = (*x*_1_, …, *x*_*L*_) of length *L* given its corrupted version 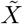 of length 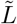 as:

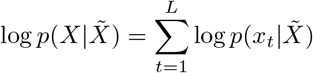

We model the joint distribution 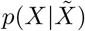 with the proposed architecture (Supplementary Fig. 2). First, we apply a sequence corruption process 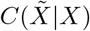 that takes as input sequence *X* of length *L* and returns corrupted version 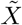, potentially of different length 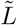 (Sec. A.1). The corrupted sequence 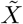 is passed as an input to the encoder 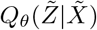 that maps 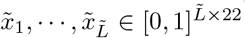 to a sequence of continuous representations 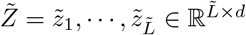. To predict the probabilities of target tokens *X* = (*x*_1_, … *x*_*L*_), we use a monotonic location-based attention mechanism [25] to transform 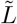 vectors of 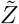 into *L* vectors. The length transform 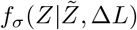 takes in 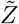 and the length difference 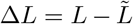 and returns the new vectors of the target sequence *Z*, with 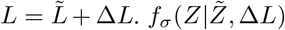 then computes each *z*_*i*_ as a weighted sum of the vectors from the encoder, i.e., 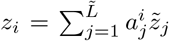 where 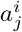 is an attention coefficient computed as in Shu *et al*. [25]. *Z* is then passed to the decoder *P*_*ϕ*_(*X|Z*), which predicts the probabilities of the target tokens.

### 3.2 Length prediction and transformation

Although the length difference is known during training, it is not readily available at inference time and must be predicted. We use an approach previously proposed by Shu *et al*. [25] and Lee *et al*. [23] and construct a length predictor as a classifier that outputs a categorical distribution 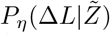 over the length difference. The classifier takes in a sequence-level representation obtained by pooling representations 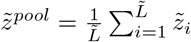, and produces a categorical distribution that covers the maximum range of length differences, [−Δ*L*_*max*_, Δ*L*_*max*_], where Δ*L*_*max*_ is determined by the choice of a corruption process (Supplementary Material, Sec. A.1). The classifier is parameterized by a single, fully connected linear layer with a *softmax* output.

### 3.3 A protein function classifier

For conditional sequence design, we incorporate a protein function classifier by training it on the representations from the encoder of the DAE. The goal is to exploit the error signal from the function classifier at test time to guide sampling towards sequences with desired functions. We train a multi-label classifier 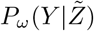 that takes in a latent representation 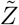 from the encoder and that outputs a vector *Y* of probabilities for each function as well as the classifier’s internal latent representation *Z*_*c*_. We parameterize *P*_*ω*_ with one multi-head attention (MHA) layer that maps the initial sequence feature representation 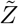 to an internal feature representation, 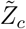 of the same hidden dimension as 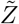, which is pooled to form a protein-level representation; 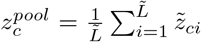. This protein-level representation is passed to single, fully connected layer followed by a point-wise sigmoid function that returns function probabilities.

### 3.4 Function-conditioned sampling

We guide sampling towards a target function *i* at every sampling step by using the gradient of the function classifier’s predictive probability of *i* to update the encoder’s vectors and increase the likelihood of higher expression of the desired target function. At every generation step, we update the internal state of the encoder as follows:

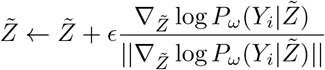

where *ϵ* controls the strength of the function gradients. The output 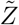 is then passed to the length transform, together with the predicted Δ*L* sampled from the length predictor, and then to the decoder (Supplementary Fig. 2).

## 4 Results

We demonstrate the applicability of our method in three different case studies: 1) adding a function to an existing fold by installing a binding site (Fig. 1A) 2) diversifying a protein sequence by preserving its function and salient residues (Fig. 1B), and 3) modifying protein function by globally changing the protein fold (Supplementary Material, Sec. A.4).

**Figure 1:**
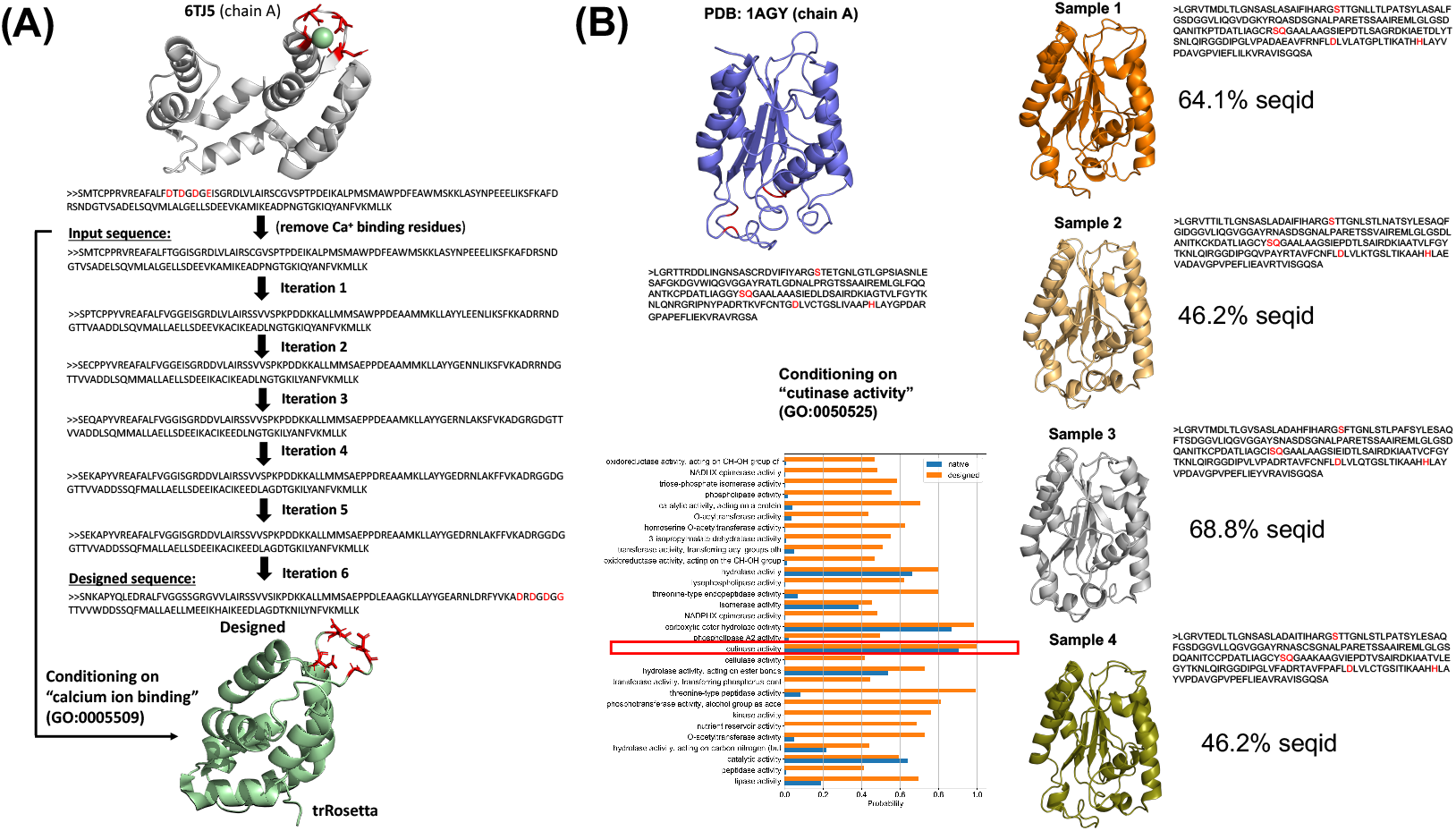
(A) A designed sequence of 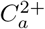-binding protein obtained by altering the sequence of calmodulin, calcium-binding protein (PDB: 6TJ5, chain A) after removing its calcium binding site. (B) Redesign of *fusarium solani pisi* cutinase (PDB ID: 1AGY, chain A) cutinases with enhanced functions.

### Designing a sequence with a metal binding site

We study the ability of the model to add a potential metal binding sites to a protein. In particular, we test the model’s ability to recover metal binding sites by starting the sampling procedure from a sequence of a metal binding protein after removing the known binding residues, including residues involved in calcium binding (three aspartate and one glutamic amino acid residues) from a calcium-binding protein (PDB: 6TJ5, chain A; Fig. 1A). Starting from the altered sequence, we perform sampling by conditioning on *calcium ion binding (GO:0005509)*. After six MCMC steps, we obtain a sequence with a high score for calcium ion binding and observe a sequence motif frequently found in most known calcium binding proteins (Fig. 1A); highlighted in red). The designed sequence has 48.7% sequence identity to the the starting one. When folded using the *trRosetta* package [30], it forms a helix-loop-helix structural domain at the location of the predicted binding site [31]. The three aspartite amino acids in this loop are negatively charged and interact with a positively charged calcium ion. The glycine is necessarily due to the conformational requirements of the backbone [31, 32].

### Redesign of cutinases with enhanced functions

We test the ability of model to diversify an existing protein sequence by preserving the functional residues. Here, we use sequence of *fusarium solani pisi* cutinase. Cutinases are responsible for hydrolysis of the protective cutin lipid layer in plants and thus have been used for hydrolysis of small molecule esters and polyesters. Sampled sequences obtained after 6 generations of sampling steps with the constrain imposed on *cutinase activity (GO:0050525)* are folded by trRosetta. The results are show in Fig.1B with catalytic residues are highlighted in red. Our function classifier shows the probability scores for *cutinase activity* of the designed sequences. We perform multiple sequence alignment of the top scoring sampled sequences showing the catalytic residues of the initial cutinase (1AGY-A) preserved by our manifold sampling strategy.

## A Supplementary Material

### A.1 Sequence corruption process

The noise-corruption processes, 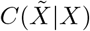, is modeled by perturbing the original sequence by applying one of three procedures:

- Removing Δ*L* residues from randomly chosen positions in the input sequence
- Inserting Δ*L* randomly chosen residues at randomly chosen positions in the input sequence resulting in a sequence with
- Mutating Δ*L* randomly chosen residues

where length difference is randomly chosen from a predefined range, Δ*L* ∈ [−Δ*L*_*max*_, Δ*L*_*max*_].

### A.2 Data collection

Unsupervised training of our model is done using ∼20M sequences from the protein family database, Pfam. Sequences longer than 1000 residues and shorter than 50 residues are removed from our training set. The dataset is randomly partitioned into training and validation sets using an 80:20 ratio. Supervised training of the function predictor is done using protein sequences with annotations for at least one Molecular Function Gene Ontology (GO) term from the Uniprot database [33]. Only GO terms with at least 50 training examples are considered.

### A.3 Training

We use a Transformer-like architecture [34] to model the encoder and decoder using a stacked MHA layer and point-wise, fully connected layers with residual connections followed by a layer normalization.

During training, both perturbed input *x* and target sequence *y* are given to the model, and the model is aware of their lengths. However, during inference the length of the target sequence has to be predicted first. Given a sequence-level embedding vector, *z*_*pool*_ we train the model to predict the length difference between *l*_*y*_ and *l*_*x*_. In our implementation, *p*(*l*_*y*_ *− l*_*x*_ | *z*_*pool*_) is modeled as a *softmax* probability distribution that covers the length difference [−Δ*L*_*max*_, Δ*L*_*max*_]. During inference, the length of the target sequence automatically adapts itself by first predicting the length difference and then by transforming the input sequence into the target sequence using the upsampling step presented in Section 3.2.

#### Algorithm 1: Manifold sampler

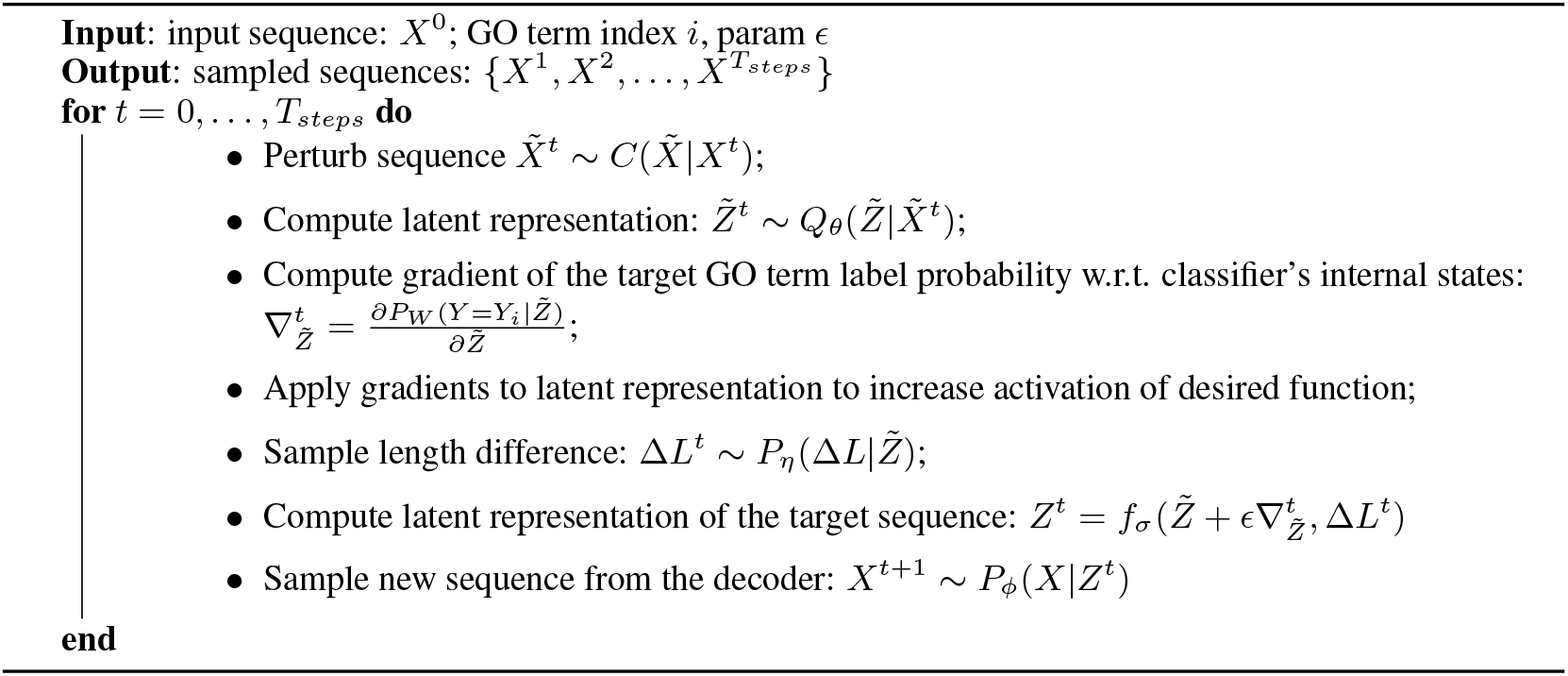

### A.4 Designing protein sequences with novel secondary structures

We explore the possibility of designing a protein with an *all-alpha* fold – a secondary structure that is almost exclusively *α*-helical – starting from a sequence of a protein having *all-beta* fold, with a secondary structure composed almost exclusively of *β*-sheets). We started with Beta-2-microglobulin protein (PDB: 4N0F, chain B) which is composed only of beta-sheets. We perform the sampling by conditioning on *ion transmembrane transporter activity (GO:0015075). α*-helices are the most common protein structure elements embedded in membranes, so the designed sequence is expected to be composed of *α*-helices. The sampling results after seven MCMC steps are shown in Fig. 3. The designed sequence has no known homologs in the PDB and has only maximal 36% sequence identical to the sequences in the Uniprot database. The sequence is folded using *trRosetta* package [30]. Using an external protein function classifier, we show that the designed sequence is predicted to have a desired function.

**Figure 2:**
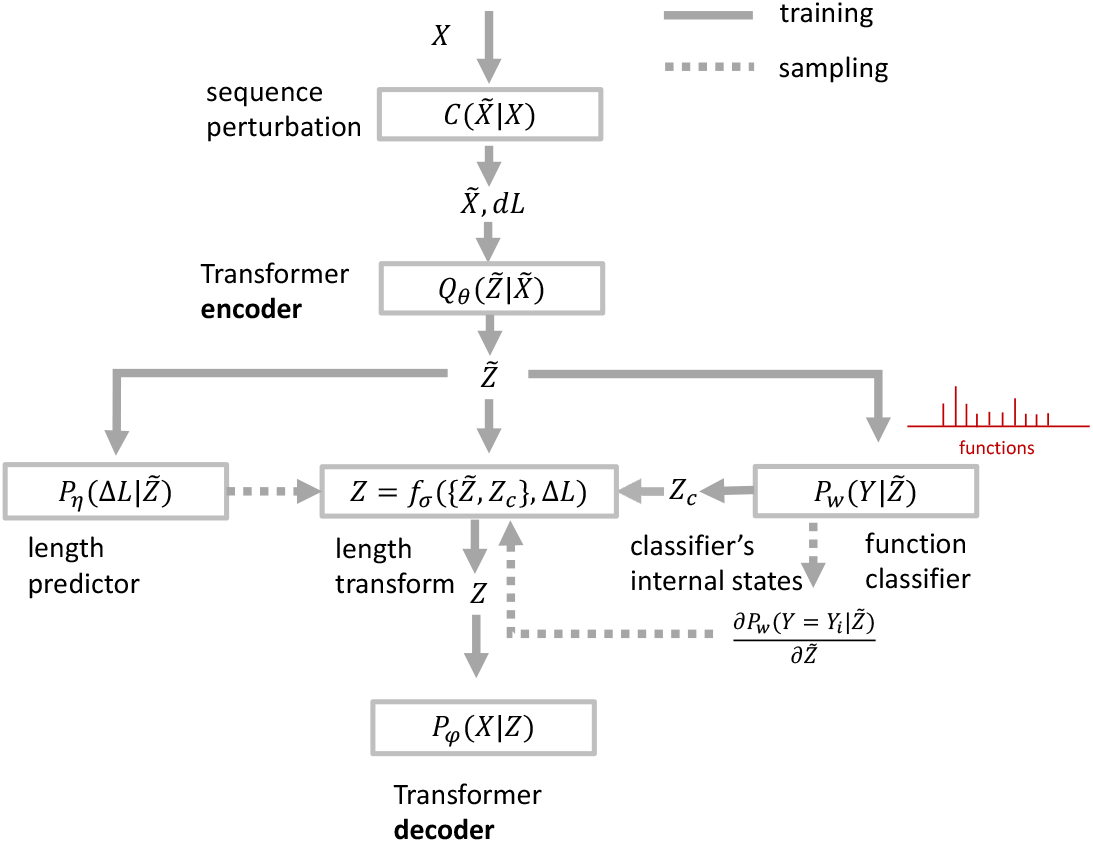
Architecture of the sequence DAE.

**Figure 3:**
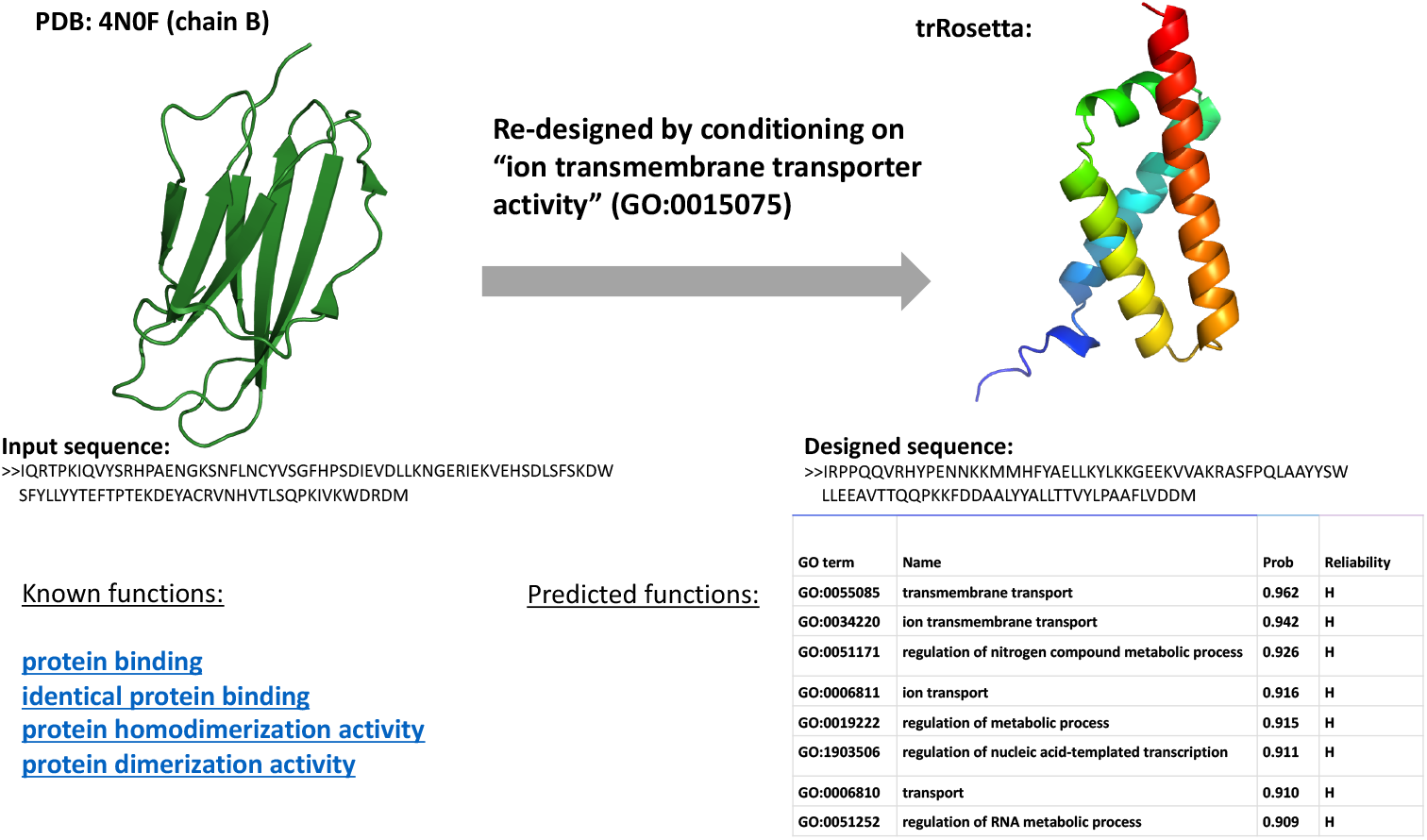
A designed sequence of *α*-helical protein obtained by altering the sequence of a *β*-protein by conditioning the sampling process with *ion transmembrane transporter activity* function label.

## References

[1] Po-Ssu Huang, Scott E. Boyken, and David Baker. The coming of age of de novo protein design. Nature, 537(7620):320–327, 2016.

[2] Philip A. Romero and Frances H. Arnold. Exploring protein fitness landscapes by directed evolution. Nature Reviews Molecular Cell Biology, 10(12):866–876, 2009.

[3] Ethan C. Alley, Grigory Khimulya, Surojit Biswas, Mohammed AlQuraishi, and George M. Church. Unified rational protein engineering with sequence-based deep representation learning. Nature Methods, 16(12):1315–1322, 2019.

[4] Tristan Bepler and Bonnie Berger. Learning protein sequence embeddings using information from structure. In International Conference on Learning Representations, 2019.

[5] Alexander Rives, Joshua Meier, Tom Sercu, Siddharth Goyal, Zeming Lin, Demi Guo, Myle Ott, C. Lawrence Zitnick, Jerry Ma, and Rob Fergus. Biological structure and function emerge from scaling unsupervised learning to 250 million protein sequences. bioRxiv, 2020.

[6] Ali Madani, Bryan McCann, Nikhil Naik, Nitish Shirish Keskar, Namrata Anand, Raphael R. Eguchi, Po-Ssu Huang, and Richard Socher. Progen: Language modeling for protein generation. bioRxiv, 2020.

[7] Roshan Rao, Nicholas Bhattacharya, Neil Thomas, Yan Duan, Peter Chen, John Canny, Pieter Abbeel, and Yun Song. Evaluating protein transfer learning with tape. In H. Wallach, H. Larochelle, A. Beygelzimer, F. Alché-Buc, E. Fox, and R. Garnett, editors, Advances in Neural Information Processing Systems 32, pages 9689–9701. Curran Associates, Inc., 2019.

[8] Jung-Eun Shin, Adam J Riesselman, Aaron W Kollasch, Conor McMahon, Elana Simon, Chris Sander, Aashish Manglik, Andrew C Kruse, and Debora S Marks.Protein design and variant prediction using autoregressive generative models. Nature communications, 12(1):1–11, 2021.

[9] Amos Bairoch and Rolf Apweiler. The SWISS-PROT protein sequence database and its supplement TrEMBL in 2000. Nucleic Acids Research, 28(1):45–48, 01 2000.

[10] Tristan Bepler and Bonnie Berger. Learning protein sequence embeddings using information from structure. In International Conference on Learning Representations, 2019.

[11] Vladimir Gligorijevic, P. Douglas Renfrew, Tomasz Kosciolek, Julia Koehler Leman, Kyunghyun Cho, Tommi Vatanen, Daniel Berenberg, Bryn Taylor, Ian M. Fisk, Ramnik J. Xavier, Rob Knight, and Richard Bonneau. Structure-based function prediction using graph convolutional networks. bioRxiv, 2019.

[12] Namrata Anand and Possu Huang. Generative modeling for protein structures. In S. Bengio, H. Wallach, H. Larochelle, K. Grauman, N. Cesa-Bianchi, and R. Garnett, editors, Advances in Neural Information Processing Systems 31, pages 7494–7505. Curran Associates, Inc., 2018.

[13] Raphael R. Eguchi, Namrata Anand, Christian A. Choe, and Po-Ssu Huang. Ig-vae: Generative modeling of immunoglobulin proteins by direct 3d coordinate generation. bioRxiv, 2020.

[14] Sivaraman Balakrishnan, Hetunandan Kamisetty, Jaime G. Carbonell, Su-In Lee, and Christopher James Langmead. Learning generative models for protein fold families. Proteins: Structure, Function, and Bioinformatics, 79(4):1061–1078, 2011.

[15] Alex Hawkins-Hooker, Florence Depardieu, Sebastien Baur, Guillaume Couairon, Arthur Chen, and David Bikard. Generating functional protein variants with variational autoencoders. BioRxiv, 2020.

[16] Kristian Davidsen, Branden J Olson, III DeWitt, William S, Jean Feng, Elias Harkins, Philip Bradley, and IV Matsen, Frederick A. Deep generative models for t cell receptor protein sequences. eLife, 8:e46935, 2019.

[17] Joe G. Greener, Lewis Moffat, and David T. Jones. Design of metalloproteins and novel protein folds using variational autoencoders. Scientific Reports, 8(1):16189, 2018.

[18] Pascal Vincent, Hugo Larochelle, Yoshua Bengio, and Pierre-Antoine Manzagol. Extracting and composing robust features with denoising autoencoders. In Proceedings of the Twenty-fifth International Conference on Machine Learning (ICML’08), pages 1096–1103. ACM, 2008.

[19] Pascal Vincent, Hugo Larochelle, Isabelle Lajoie, Yoshua Bengio, Pierre-Antoine Manzagol, and Léon Bottou. Stacked denoising autoencoders: Learning useful representations in a deep network with a local denoising criterion. Journal of machine learning research, 11(12), 2010.

[20] Pascal Vincent. A connection between score matching and denoising autoencoders. Neural computation, 23(7):1661–1674, 2011.

[21] Yoshua Bengio, Li Yao, Guillaume Alain, and Pascal Vincent. Generalized denoising auto-encoders as generative models. In Proceedings of the 26th International Conference on Neural Information Processing Systems - Volume 1, NIPS’13, page 899–907, Red Hook, NY, USA, 2013. Curran Associates Inc.

[22] Kyunghyun Cho. Noisy parallel approximate decoding for conditional recurrent language model, 2016.

[23] Jason Lee, Elman Mansimov, and Kyunghyun Cho. Deterministic non-autoregressive neural sequence modeling by iterative refinement. In Proceedings of the 2018 Conference on Empirical Methods in Natural Language Processing, 2018.

[24] Jiatao Gu, James Bradbury, Caiming Xiong, Victor O.K. Li, and Richard Socher. Non-autoregressive neural machine translation. In International Conference on Learning Representations, 2018.

[25] Raphael Shu, Jason Lee, Hideki Nakayama, and Kyunghyun Cho. Latent-variable non-autoregressive neural machine translation with deterministic inference using a delta posterior. AAAI, 2020.

[26] Jason Lee, Dustin Tran, Orhan Firat, and Kyunghyun Cho. On the discrepancy between density estimation and sequence generation. arXiv preprint 2002.07233, 2020.

[27] Samy Bengio, Oriol Vinyals, Navdeep Jaitly, and Noam Shazeer. Scheduled sampling for sequence prediction with recurrent neural networks. In Proceedings of the 28th International Conference on Neural Information Processing Systems - Volume 1, NIPS’15, page 1171–1179, Cambridge, MA, USA, 2015. MIT Press.

[28] Marc’Aurelio Ranzato, Sumit Chopra, Michael Auli, and Wojciech Zaremba. Sequence level training with recurrent neural networks, 2016.

[29] Felix Hill, Kyunghyun Cho, and Anna Korhonen. Learning distributed representations of sentences from unlabelled data. CoRR, abs/1602.03483, 2016.

[30] Jianyi Yang, Ivan Anishchenko, Hahnbeom Park, Zhenling Peng, Sergey Ovchinnikov, and David Baker. Improved protein structure prediction using predicted interresidue orientations. Proceedings of the National Academy of Sciences, 117(3):1496–1503, 2020.

[31] Jessica L. Gifford, Michael P. Walsh, and Hans J. Vogel. Structures and metal-ion-binding properties of the Ca2+-binding helix–loop–helix EF-hand motifs. Biochemical Journal, 405(2):199–221, 06 2007.

[32] Todor Dudev and Carmay Lim. Competition among metal ions for protein binding sites: Determinants of metal ion selectivity in proteins. Chemical Reviews, 114(1):538–556, 01 2014.

[33] Rolf Apweiler, Amos Bairoch, Cathy H. Wu, Winona C. Barker, Brigitte Boeckmann, Serenella Ferro, Elisabeth Gasteiger, Hongzhan Huang, Rodrigo Lopez, Michele Magrane, Maria J. Martin, Darren A. Natale, Claire O’Donovan, Nicole Redaschi, and Lai-Su L. Yeh. Uniprot: the universal protein knowledgebase. NUCLEIC ACIDS RES, 32:115–119, 2004.

[34] Ashish Vaswani, Noam Shazeer, Niki Parmar, Jakob Uszkoreit, Llion Jones, Aidan N. Gomez, undefine-dukasz Kaiser, and Illia Polosukhin. Attention is all you need. In Proceedings of the 31st International Conference on Neural Information Processing Systems, NIPS’17, page 6000–6010, Red Hook, NY, USA, 2017. Curran Associates Inc.

